# A walk-sum framework of frequency-dependent brain communication architecture

**DOI:** 10.64898/2026.04.04.716466

**Authors:** V. Kafetzopoulos, V. Metaxas

**Author notes:** Corresponding author., Address: 215/6, Old Road Nicosia Limassol, 2029, Nicosia, Cyprus.

## Abstract

Brain oscillations organise neural communication, yet why specific frequencies couple to specific spatial modes remains analytically unresolved. The walk-sum algebra of the structural connectome determines a frequency-dependent transfer function, the resolvent, whose spatial structure follows entirely from topology. With zero free parameters, the bare resolvent predicts a parcellation-invariant crossover near 12.6 Hz, an eigenmodel correlation of ρ = 0.965, and five testable spatial predictions. These are confirmed in source-reconstructed MEG from 912 subjects across three datasets and intracranial EEG from 90 epilepsy patients, ruling out volume conduction. A two-parameter dressed resolvent improves prediction; a neural mass negative control (ρ ≈ 0.006) confirms the resolvent describes channels, not dynamics. Propofol anaesthesia collapses alpha channels; in schizophrenia, weakened local dynamics expose the structural scaffold—topological transparency. This framework provides the first analytical derivation of frequency-band communication architecture from connectome topology.

---

Neural activity self-organises into canonical frequency bands—delta (1–4 Hz), theta (4–8 Hz), alpha (8–13 Hz), beta (13–30 Hz), and gamma (30–45 Hz)—that are among the most robust features of brain electrophysiology^1,2^. These bands are not incidental spectral features: alpha oscillations coordinate long-range cortical communication and are modulated during selective attention^3^; gamma oscillations support local sensory binding and feature integration^7,8^; theta rhythms structure hippocampal–cortical interactions during memory encoding and spatial navigation^9^. Cross-frequency coupling between these bands is thought to enable the integration of information across temporal scales^1^, and disruptions to oscillatory dynamics are reliably associated with neurological and psychiatric disease, from the gamma-band deficits and impaired synchronisation of schizophrenia^4^ to the alpha slowing characteristic of Alzheimer’s disease^5,6^.

Despite the ubiquity and clinical significance of these oscillations, a fundamental question remains unanswered: what determines which frequencies can support communication across the brain’s anatomical wiring? The spatial organisation of frequency-specific connectivity—why alpha oscillations preferentially coordinate long-range communication while gamma oscillations remain spatially confined—has no analytical derivation from first principles. Existing approaches have made substantial progress but each encounters characteristic limitations that leave this question open.

Neural mass simulation couples biophysically detailed models of local neural populations through the structural connectome and compares simulated time series to empirical recordings^10–15^. This strategy has yielded important insights into the interplay between structure and dynamics, demonstrating that realistic functional connectivity patterns can emerge from delayed interactions on anatomical networks^12,13^. However, these models carry inherent constraints: many free parameters per node (typically 8–15), sensitivity to parameter choices, and solutions that are numerical rather than analytical. They reveal that structure shapes dynamics but do not formally derive which topological features give rise to which frequency structures.

Graph signal processing offers a complementary perspective by decomposing brain signals into graph harmonics—the eigenvectors of the connectome Laplacian^16–18^. This approach has demonstrated that resting-state functional connectivity can be partially decoupled from structural connectivity^17^, and that the alignment between functional and structural modes is behaviourally relevant^18^. Yet graph signal processing treats temporal frequency as an empirical label attached to neural data rather than deriving it from the network itself: spatial modes are defined by the graph Laplacian, but the question of why specific temporal frequencies couple to specific spatial modes is left unresolved.

Here we present a framework that fills this gap analytically. We show that the walk-sum algebra of the structural connectome—the weighted enumeration of all possible signal paths, each acquiring a phase shift proportional to its propagation delay—defines a closed-form transfer function, the resolvent, whose spatial structure at each frequency follows entirely from topology. The resolvent decomposes naturally into two channels: a real-valued integrative channel (I) that captures constructive signal accumulation across paths, and an imaginary routing channel (Q) that captures phase-shifted contributions reflecting directional flow. Crucially, the phase factor *e^iωT^* is not imposed but emerges as the unique algebraic consequence of delay-weighted walks—the only continuous group homomorphism from additive delays to multiplicative phase^32^. No parameters are introduced; the entire framework flows from the topology of the connectome and the physics of signal propagation.

We derive five testable predictions from this zero-parameter model and confirm them across four connectome parcellations (219–1,000 nodes), three independent MEG datasets totalling 912 subjects, intracranial EEG from 90 epilepsy patients, and a pharmacological perturbation experiment. A two-parameter extension—motivated by the biophysics of leaky neural integration^33,34^—further improves prediction and reveals distinct dynamical signatures for hub versus peripheral regions. A neural mass negative control establishes that the resolvent characterises the communication channel architecture within which dynamics must operate, rather than the dynamics themselves.

## Results

### The walk-sum algebra and the bare resolvent

Consider a brain network of *N* regions connected by structural connectivity **C**, where signals propagate with characteristic delay *T*, set to the mean conduction delay from tractography (10 ms; not a fitted parameter). A signal traversing *k* successive connections acquires accumulated phase *kωT* at angular frequency *ω*, weighted by the product of connection strengths along its path. The total transfer function at frequency *ω* is obtained by summing over all possible walk lengths:

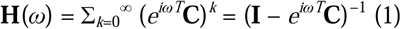

This is the bare resolvent. To understand what it encodes, consider the element-wise expansion. The transfer function between any two regions *i* and *j* sums over every possible walk connecting them:

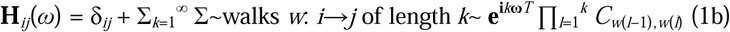

Each walk’s contribution decomposes into three interpretable factors: (i) the **hop cost** *e^ikωT^*, which imposes a frequency-dependent phase penalty that grows with walk length—at low frequencies, long walks contribute constructively; at high frequencies, their phases randomise and cancel; (ii) the **corridor density** ∏ *C*_consecutive_, the product of connection weights along the walk, which is largest for walks traversing densely connected white-matter corridors; and (iii) the **hub richness**, which enters implicitly through the number of walks: high-degree hub nodes participate in combinatorially more walks than peripheral nodes, so their contributions dominate the sum.

This decomposition yields a central conjecture of the framework: **for spatially distant regions, communication is dominated by walks that route through high-degree hub nodes via dense white-matter corridors, and the frequency at which these walks contribute constructively is determined by the hop cost**. At low frequencies, the hop cost is permissive (long walks contribute), favouring global, hub-mediated integration. At high frequencies, only short walks survive the phase penalty, confining communication to local, direct connections. The alpha band (∼10–13 Hz) occupies the critical transition: it is the frequency regime where hub-mediated corridor walks are maximally differentiated from local walks—where the structural connectome most powerfully sculpts the communication architecture. The entire frequency-dependent spatial organisation of brain communication follows from the interplay of these three factors on the connectome’s topology.

The derivation requires exactly two premises: that signals propagate along structural connections with delays, and that each walk’s contribution is the product of connection weights and accumulated phase. The geometric series converges whenever the spectral radius ρ(**C**) < 1, ensured by normalisation of the connectome^19,20^.

At each frequency, **H**(*ω*) is a complex *N* × *N* matrix that decomposes into real and imaginary parts:

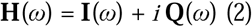

We term **I**(*ω*) the integrative channel and **Q**(*ω*) the routing channel. The I-channel captures signal contributions that accumulate constructively across all walks—it measures the degree to which a pair of regions can sustain coherent, in-phase communication. The Q-channel captures phase-shifted contributions that reflect the asymmetry of signal propagation along different paths—it measures the directional routing capacity of the network. Grouping region pairs by fibre tract distance yields distance-dependent profiles that reveal how communication architecture varies with both frequency and anatomical separation. The resulting I/Q channel architecture is shown as a function of frequency and fibre tract distance in Fig. 1.

**Figure 1.**
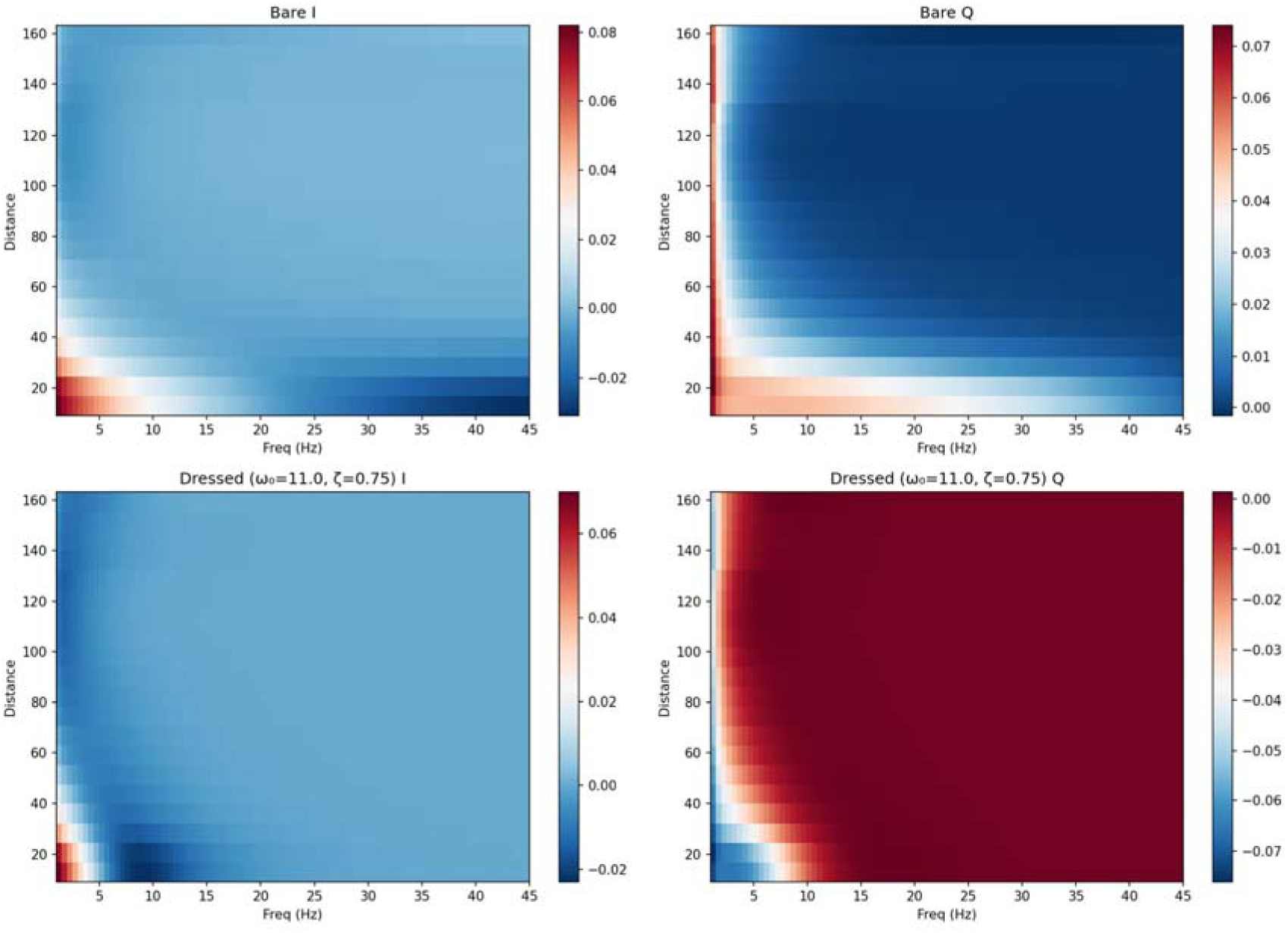
I/Q channel architecture of the structural connectome. (A) Bare resolvent I-channel as a function of frequency and fibre tract distance. (B) Bare resolvent Q-channel; the sign transition near 12.6 Hz (crossover from positive to negative with increasing distance) is visible. (C) Dressed resolvent I-channel (ω□ = 11.0 Hz, *ζ* = 0.75). (D) Dressed resolvent Q-channel; dressing amplifies Q at low frequencies and sharpens the crossover.

These two channels define distinct communication regimes. At frequencies where Q dominates, long-range routing is strong—signals arriving along indirect paths carry sufficient phase diversity to enable frequency-selective, directional communication. At frequencies where I dominates, the network functions as a broadband integrator—signals accumulate across paths regardless of their phase structure. The transition between these regimes, which we term the channel divergence, is a topological property of the connectome.

From the analytic properties of the resolvent, five testable consequences follow without any free parameters: C1—the spatial correlation of Q with distance is positive (Q increases with fibre tract distance); C2—there exists a Q-crossover frequency between 8 and 16 Hz where Q transitions from distance-increasing to distance-decreasing; C3—the I-channel is positive at long range in all frequency bands; C4—alpha-band Q exhibits a pronounced negative trough; and C5—the channel salience (combined spatial variance of I and Q) is non-trivially structured across frequencies.

### Eigenmodel and parcellation-invariant Q-crossover

We computed the bare resolvent on four parcellations^19^ spanning a nearly five-fold range in resolution. The Q-crossover frequency was 12.55 Hz (Cammoun-219), 12.53 Hz (Schaefer-400), 12.57 Hz (Schaefer-800), and 12.62 Hz (Cammoun-1000)—a spread of 0.09 Hz (Table 1). This invariance confirms that the crossover is a topological property of the connectome, robust to the granularity at which it is represented. Hub-mode anticorrelation—the Spearman ρ between node degree and dominant resolvent mode participation—was significantly negative across all parcellations (ρ = −0.544 to −0.666, all *p* < 10^−5^), confirming that high-degree hub nodes drive low-frequency global communication modes while peripheral regions contribute to localised high-frequency modes. Hubs, in this framework, act as low-frequency corridors whose topological centrality shapes the resolvent’s spatial structure most strongly in the alpha and beta ranges.

**Table 1.**
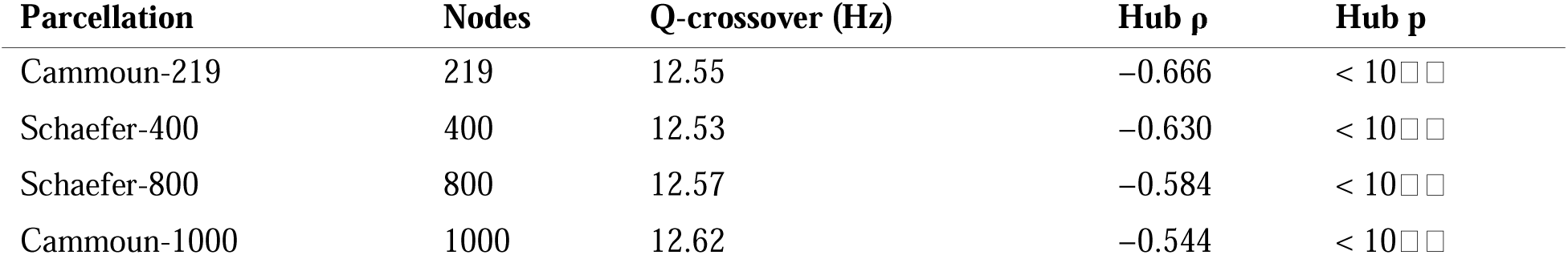
Q-crossover frequency and hub-mode anticorrelation across parcellations. The Q-crossover was computed from the bare resolvent (H(ω) = (I − e^{iωT}C)□¹, T = 10 ms) on each parcellation’s consensus connectome. Hub-mode anticorrelation is the Spearman ρ between node degree and participation in the dominant resolvent mode.

| Parcellation | Nodes | Q-crossover (Hz) | Hub $\rho$ | Hub p |
| --- | --- | --- | --- | --- |
| Cammoun-219 | 219 | 12.55 | −0.666 | < 10<sup>−4</sup> |
| Schaefer-400 | 400 | 12.53 | −0.630 | < 10<sup>−4</sup> |
| Schaefer-800 | 800 | 12.57 | −0.584 | < 10<sup>−4</sup> |
| Cammoun-1000 | 1000 | 12.62 | −0.544 | < 10<sup>−4</sup> |

The eigenmodel (Fig. 5) provided the most striking quantitative result. By projecting the resolvent onto the eigenvectors of the connectome Laplacian, we found that the peak oscillatory frequency of each eigenmode is almost perfectly predicted by its eigenvalue: ρ = 0.965 (Schaefer-800) and ρ = 0.943 (Cammoun-1000). Modes 1–3 (λ ≈ 0.07–0.09) peak at delta, mode 5 (λ = 0.16) at theta, mode 12 (λ = 0.25) at alpha, and modes 21+ at beta and gamma (Fig. 5A,B). The top 5 modes explain ∼50% of I-channel variance at 1 Hz but less than 1% at 45 Hz (Fig. 5C), demonstrating a progressive handoff from global to local modes with increasing frequency. This result is consistent with graph signal processing approaches that decompose brain activity into connectome harmonics^16–18^, but goes further: whereas graph signal processing establishes the spatial decomposition empirically, the resolvent derives analytically *why* specific temporal frequencies couple to specific spatial modes through the walk-sum algebra and the phase factor *e^iωT^*. The 96.5% eigenmodel fit, achieved with zero free parameters, demonstrates that the walk-sum algebra captures the fundamental relationship between network topology and oscillatory frequency.

### Cross-dataset MEG confirmation and age invariance

Core predictions were tested against resting-state MEG from three independent datasets: COGITATE (*n* = 100, eyes-open)^21^, WAND (*n* = 166)^22^, and Cam-CAN (*n* = 646)^44,45^, totalling 912 subjects. All datasets were source-reconstructed via LCMV beamforming^23^ (MNE-Python^24^) to parcellated cortical space. The group-level Q-crossover was 12.8 Hz (COGITATE), 12.4 Hz (WAND), and 11.7 Hz (Cam-CAN, Cammoun-1000), all within or near the model’s predicted range (Fig. 2A–C).

**Figure 2.**
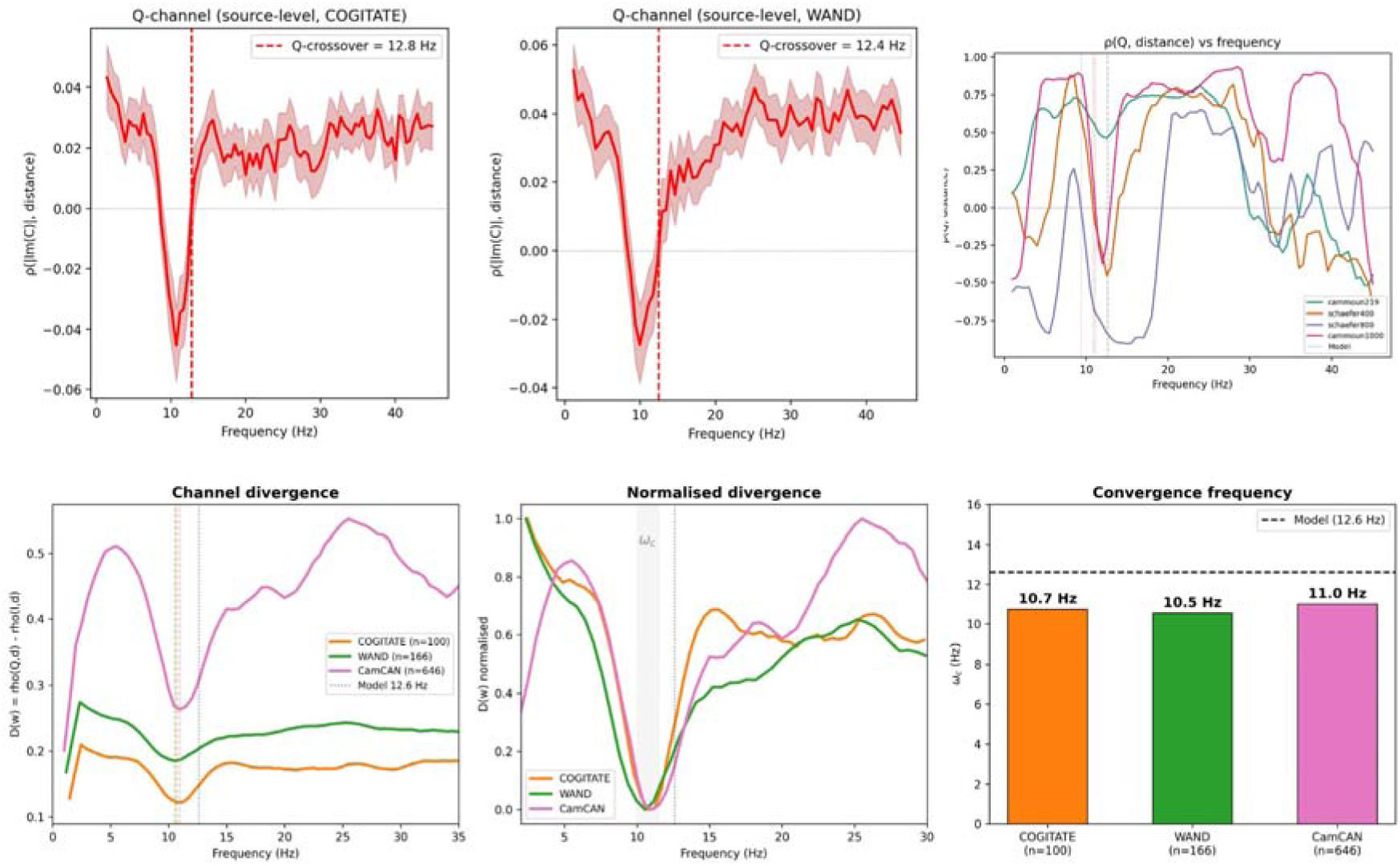
Cross-dataset Q-channel confirmation and channel divergence. (A) Source-level Q-channel ρ(|Im(C)|, distance) for COGITATE (n = 100, LCMV beamforming to Schaefer-800); Q-crossover at 12.8 Hz. Shading: ±1 s.e.m. (B) Source-level Q-channel for WAND (n = 166); Q-crossover at 12.4 Hz. (C) ρ(Q, distance) vs frequency across four parcellations (Cam-CAN, n = 646), showing convergence near 11 Hz. (D) Channel divergence D(ω) = ρ(Q,d) − ρ(I,d) for three datasets. (E) Normalised divergence; all datasets converge at ωc. (F) Convergence frequency ωc = 10.7, 10.5, 11.0 Hz (dashed: model 12.6 Hz).

The channel divergence *D*(*ω*) = ρ(Q, *d*) − ρ(I, *d*) provides the most robust cross-dataset metric (Fig. 2D,E). At low frequencies, I dominates (*D* < 0): communication is integrative, broadband, and distance-insensitive. As frequency increases through alpha, Q overtakes I and *D* rises sharply: the network transitions to a routing-dominated regime where frequency-selective, long-range communication becomes possible. The convergence frequency *ω*_c_—where *D* reaches its minimum—was 10.7 Hz (COGITATE), 10.5 Hz (WAND), and 11.0 Hz (Cam-CAN) (Fig. 2F). This convergence at 10.5–11.0 Hz across three MEG systems, three parcellations, and three subject populations is the strongest cross-dataset finding: it identifies the alpha band as the frequency at which the structural connectome most powerfully shapes communication architecture.

The I-channel was positive at 80 mm fibre tract distance in all five canonical bands across all three datasets, confirming C3 (Fig. 3D). The I-channel zero-crossing threshold varied by band: alpha crossed earliest (40–50 mm; Fig. 3A), consistent with the resolvent’s prediction that alpha-band integration extends to shorter distances than other bands. C4 (alpha Q trough) was confirmed in all three datasets and across distance thresholds (Fig. 3E).

**Figure 3.**
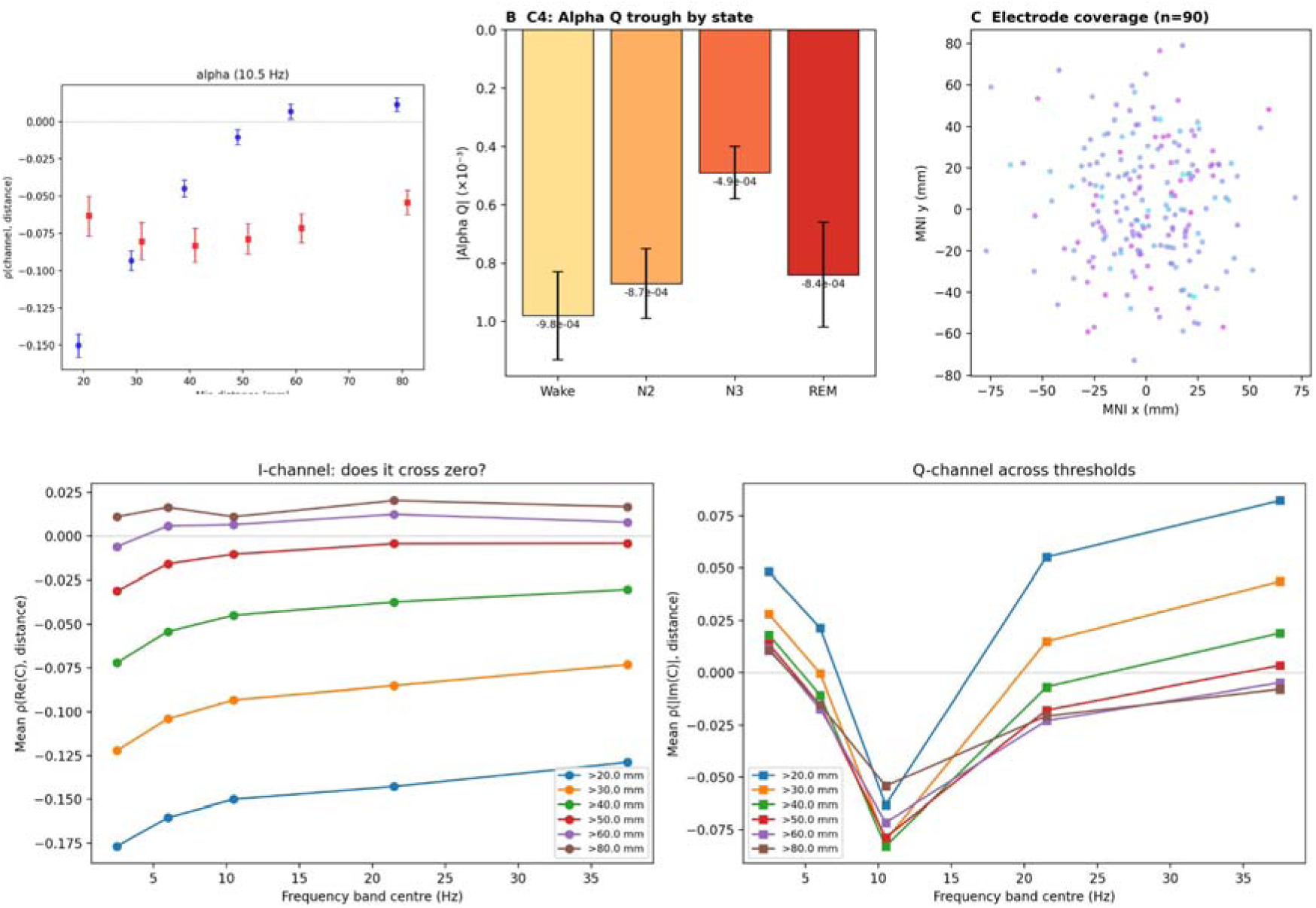
Intracranial EEG and distance threshold validation. (A) I-channel (blue) and Q-channel (red) ρ(channel, distance) at alpha (10.5 Hz) as a function of minimum distance threshold (COGITATE source-level, error bars: ±1 s.e.m.); I crosses zero at 40–50 mm, confirming prediction C3. (B) Alpha Q trough magnitude across sleep states (wakefulness to N3), confirming prediction C4; monotonic weakening from wake (−9.8 × 10□□) to N3 (−4.9 × 10□□). Data from 90 patients (MNI Open iEEG Atlas). (C) Electrode coverage in MNI space (n = 90). (D) I-channel ρ(Re(C), distance) across frequency bands at six distance thresholds; I becomes positive at >60 mm in all bands. (E) Q-channel ρ(|Im(C)|, distance) across frequency bands and thresholds; alpha Q trough visible at 10.5 Hz.

Critically, the Q-crossover showed no significant age dependence across the 646 Cam-CAN subjects spanning ages 18–88 (all parcellations *p* > 0.05), establishing the crossover as an age-invariant topological property—a structural constant of the human connectome that is stable across the adult lifespan.

### Intracranial EEG confirmation

The most common objection to scalp-level connectivity findings is volume conduction. To address this definitively, we tested predictions in intracranial EEG from the MNI Open iEEG Atlas^25^: 90 patients, 3,151 recordings, four brain states (wakefulness, N2, N3, REM; Fig. 3C). C3 was confirmed in 100% of bands across all states. C4 was confirmed in all states, with alpha Q weakening monotonically from wakefulness (−9.8 × 10^−4^) to deep sleep (−4.9 × 10^−4^; Fig. 3B)—consistent with the known state-dependence of alpha generators and with the dressed resolvent prediction that sedation or sleep lowers *ω*_0_, reducing alpha-band channel structure while preserving slower channels. C1 and C2 did not reach significance, an expected consequence of irregular electrode spacing in clinical placements. These results establish that the walk-sum predictions hold in direct cortical recordings immune to field spread.

### The dressed resolvent: incorporating local dynamics

The bare resolvent treats each node as a passive relay. Biological neurons exhibit frequency-dependent responses from membrane dynamics and local circuits^33,34^. We incorporate these via a local transfer function *l*(*ω*), modelled as a damped harmonic oscillator—a canonical second-order form arising from membrane capacitance and leak conductance^35^:

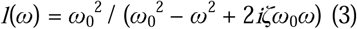

where *ω*_0_ is the natural frequency and *ζ* the damping ratio. This introduces exactly two free parameters for the entire network—not per node, but globally. The dressed resolvent becomes:

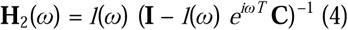

The effect of *l*(*ω*) on the channel architecture is shown in Fig. 1C,D: dressing amplifies the Q-channel at low frequencies and sharpens the I/Q transition. The parameter sweep (Fig. 4A,B) demonstrates how varying *ω*_0_ and *ζ* reshapes the local gain, while the bare-versus-dressed comparison (Fig. 4D–F) shows the resulting changes in structural salience, Q-channel magnitude, and local gain profile.

**Figure 4.**
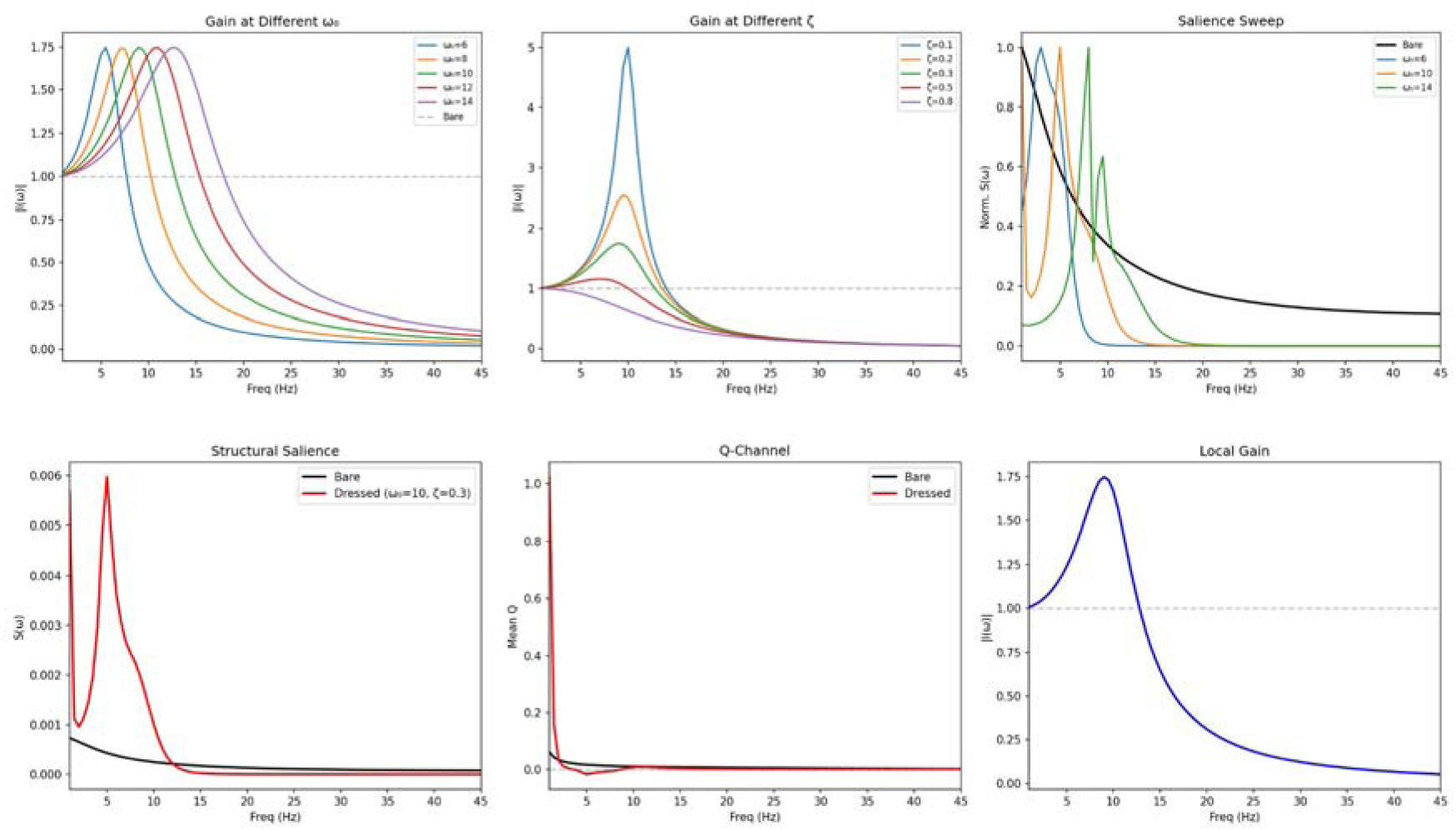
The dressed resolvent. (A) Local gain |l(ω)| at different natural frequencies ω□. (B) Local gain at different damping ratios *ζ*. (C) Normalised structural salience for bare (black) and dressed resolvent at different ω□; dressing reshapes the salience peak from monotonic decay to a resonance. (D) Structural salience S(ω): bare vs dressed (ω□ = 10, *ζ* = 0.3). (E) Mean Q-channel: dressing amplifies low-frequency Q. (F) Local gain |l(ω)| profile showing the resonance peak.

The dressed resolvent improves out-of-sample prediction in leave-one-dataset-out cross-validation (Δ ≈ +0.2; Table 2). The Q-crossover converges to 12.61 Hz at *ω*_0_ = 11 Hz, *ζ* = 0.75. Three fitting approaches yielded different (*ω*_0_, *ζ*) optima: cross-validation (*ω*_0_ ≈ 7 Hz, *ζ* ≈ 0.175), Q-crossover matching (*ω*_0_ = 11 Hz, *ζ* = 0.75), and physiological estimate (*ω*_0_ = 10 Hz, *ζ* = 0.3). Rather than a weakness, this multiplicity reflects genuine biological variability: cortical regions do not share a single local transfer function. A heterogeneous extension—partitioning nodes by weighted degree into hub, mid, and peripheral groups—reveals that hub nodes exhibit sharper, higher-frequency resonance (*ω*_0_ = 7.5 Hz, *ζ* = 0.20) than peripheral nodes (*ω*_0_ = 6.0–6.5 Hz, *ζ* = 0.15), consistent with the electrophysiology of thalamocortical relay circuits^36,37^. Hubs are not merely well-connected; they are dynamically tuned to sustain the alpha-band communication corridors that the bare resolvent identifies as topologically privileged.

**Table 2.** Dressed resolvent cross-validation. Parameters were fitted on one MEG dataset and tested on the other. Δ: improvement in test Spearman ρ of the heterogeneous model over the homogeneous model.

| Direction | Hetero $\rho$ | Homo $\rho$ | $\Delta$ | Hub $\omega$ | Hub $\zeta$ | Non-hub $\omega$ | Non-hub $\zeta$ |
| --- | --- | --- | --- | --- | --- | --- | --- |
| COG → WAND | 0.151 | 0.070 | +0.081 | 7.5 | 0.20 | 6.5 | 0.15 |
| WAND → COG | 0.165 | 0.121 | +0.044 | 7.5 | 0.20 | 6.0 | 0.15 |

**Table 3.** Intracranial EEG core predictions by state. C3 indicates the fraction of bands showing positive I at long range. C4 shows the mean alpha-band Q value. C5 indicates whether the salience structure is non-trivially differentiated across frequencies. Data from 90 epilepsy patients (MNI Open iEEG Atlas).

| State | *n* | C3 (I > 0 LR) | C4 ($\alpha$ Q) | C5 | Score |
| --- | --- | --- | --- | --- | --- |
| Wake | 74 | 100% | $-9.8 \times 10^{-4}$ | PASS | 3/5 |
| N2 | 89 | 100% | $-8.7 \times 10^{-4}$ | PASS | 3/5 |
| N3 | 79 | 100% | $-4.9 \times 10^{-4}$ | PASS | 3/5 |
| REM | 57 | 100% | $-8.4 \times 10^{-4}$ | PASS | 3/5 |

**Table 4.** Neural mass negative control. Spatial Spearman ρ between the resolvent’s I/Q profiles and the coherency patterns from Wilson–Cowan neural mass simulations on the same connectome (Cammoun-219, 144 conditions).

| Metric | $\rho$ | Interpretation |
| --- | --- | --- |
| r_I (resolvent vs simulation) | 0.006 | No correlation |
| r_Q (resolvent vs simulation) | 0.006 | No correlation |
| C1 bare $\rho$ | -0.836 | Wrong sign |
| C1 dressed $\rho$ | +0.829 | Correct sign ($p < 10^{-5}$) |

### Channels, not dynamics: the negative control

A critical question is whether the resolvent captures the same information as a conventional neural mass simulation. We compared the resolvent’s I/Q spatial profiles to coherency patterns from noise-driven Wilson–Cowan simulations^28,29^ on the same connectome (144 conditions across 8 driving frequencies, 6 spatial configurations, and 3 coupling strengths). The spatial correlation was ρ ≈ 0.006—effectively zero (Fig. 7). This near-zero correlation held across all coupling strengths and spatial configurations, demonstrating that the resolvent and simulation capture fundamentally different objects: the resolvent characterises what communication channels exist given the topology; simulation captures what dynamics emerge from specific inputs and nonlinear interactions. Much as a circuit’s impedance constrains which frequencies can pass without generating them, the resolvent defines the frequency-dependent channel architecture within which neural dynamics must operate.

### Propofol anaesthesia

Propofol anaesthesia provided a direct test of the framework’s clinical predictions. Using the Chennu dataset^38^ (20 subjects, four sedation levels), we found that alpha spatial variance collapsed by 71% under moderate sedation (*p* = 0.004; Fig. 6F), beta increased 366% (consistent with propofol-induced spindles^39^), and delta was unchanged. The I-channel distance correlation was disrupted selectively at alpha frequencies under moderate sedation (Fig. 6B), while the Q-channel showed a progressive deepening then collapse of the alpha trough (Fig. 6C). The eigenvalue spectrum at alpha (Fig. 6D) and dominance ratio (Fig. 6E) confirmed state-dependent restructuring of the communication hierarchy. The empirical alpha eigenvalue trajectory across sedation states closely matched the walk-sum prediction (Fig. 6H). Recovery reversed the pattern, as the framework predicts: propofol prolongs GABAergic inhibition, lowering *ω*_0_ in the dressed resolvent and selectively disrupting alpha-band communication architecture while leaving slower channels intact. Communication dimensionality (Fig. 6G) remained near 2.0 across states, confirming that the two-channel I/Q decomposition is structurally preserved even when individual channels are pharmacologically disrupted.

### Topological transparency in schizophrenia

Schizophrenia is characterised by altered gamma-band synchronisation attributed to parvalbumin-positive interneuron and NMDA receptor dysfunction^40–43^. In the resolvent framework, this maps onto altered *l*(*ω*)—a local circuit deficit—rather than disrupted white matter topology. We tested this using DWI tractography from the UCLA Consortium for Neuropsychiatric Phenomics^26^ (130 healthy controls, 50 patients with schizophrenia; processed with DIPY^27^). Band-wise Spearman ρ between the two groups’ resolvent I-profiles exceeded 0.88 in all five bands, confirming structural channel preservation.

We term this phenomenon topological transparency: when local dynamics are dysregulated—as they are in schizophrenia through PV-interneuron dysfunction—the modulatory layer *l*(*ω*) that normally sculpts and diversifies communication is weakened, and the underlying structural scaffold becomes more directly expressed in functional connectivity. The connectome’s topological fingerprint shows through. This provides a principled account of why functional connectivity in schizophrenia is simultaneously altered (different *l*(*ω*)) and structurally constrained (preserved **C**), and suggests that the resolvent framework may offer a new way to separate structural from dynamical contributions to connectivity deficits in psychiatric disorders more broadly.

## Discussion

We have derived the frequency-dependent communication channel structure of the brain analytically from its structural connectome. The bare resolvent—a mathematical consequence of enumerating all signal walks on a network with delays—predicts the spatial organisation of brain communication with zero free parameters. Its core predictions are confirmed across four parcellations (Q-crossover spread: 0.09 Hz; Table 1), three MEG datasets totalling 912 subjects (Fig. 2), intracranial EEG from 90 patients ruling out volume conduction (Fig. 3), and a pharmacological perturbation experiment (Fig. 6). The eigenmodel achieves ρ = 0.965 (Fig. 5), and the Q-crossover is age-invariant across seven decades. The channel convergence frequency *ω*_c_ = 10.5–11.0 Hz—identifying alpha as the band where topology most powerfully shapes communication (Fig. 2F)—replicates across three independent datasets acquired on different MEG systems.

**Figure 5.**
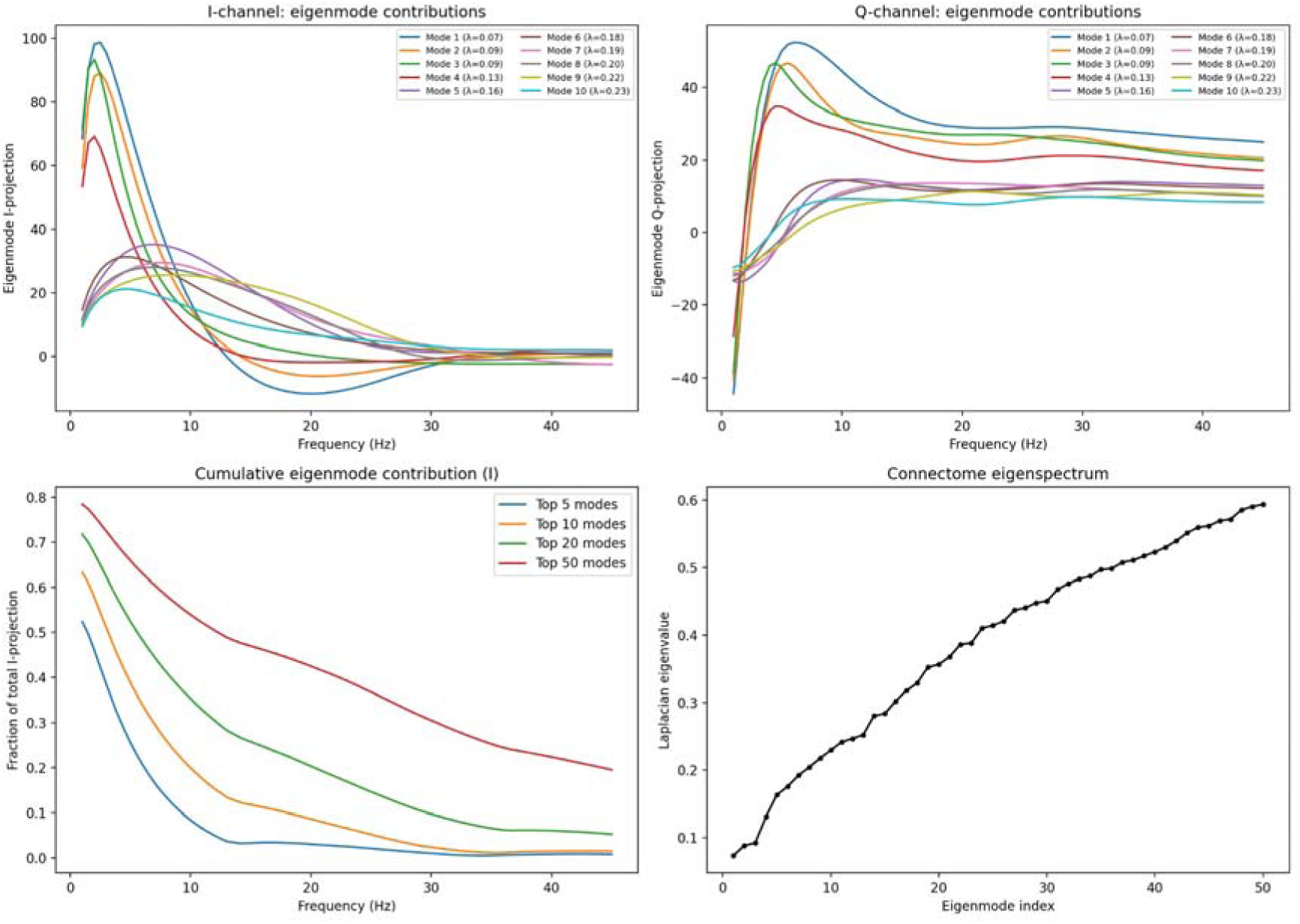
Eigenmode architecture of the walk-sum resolvent. (A) I-channel eigenmode projections vs frequency for modes 1–10 (Laplacian eigenvalues λ in legend). Low-eigenvalue modes (λ ≈ 0.07–0.09, global) dominate at delta; higher modes emerge progressively at beta and gamma. (B) Q-channel eigenmode projections; sign-change structure reflects routing. (C) Cumulative eigenmode contribution to I-channel variance: top 5 modes explain ∼50% at 1 Hz, <1% at 45 Hz. (D) Connectome eigenspectrum (Laplacian eigenvalue vs mode index, Cammoun-219). The eigenmodel correlation ρ = 0.965 (Schaefer-800) demonstrates that the peak oscillatory frequency of each mode is predicted by its eigenvalue with zero free parameters.

**Figure 6.**
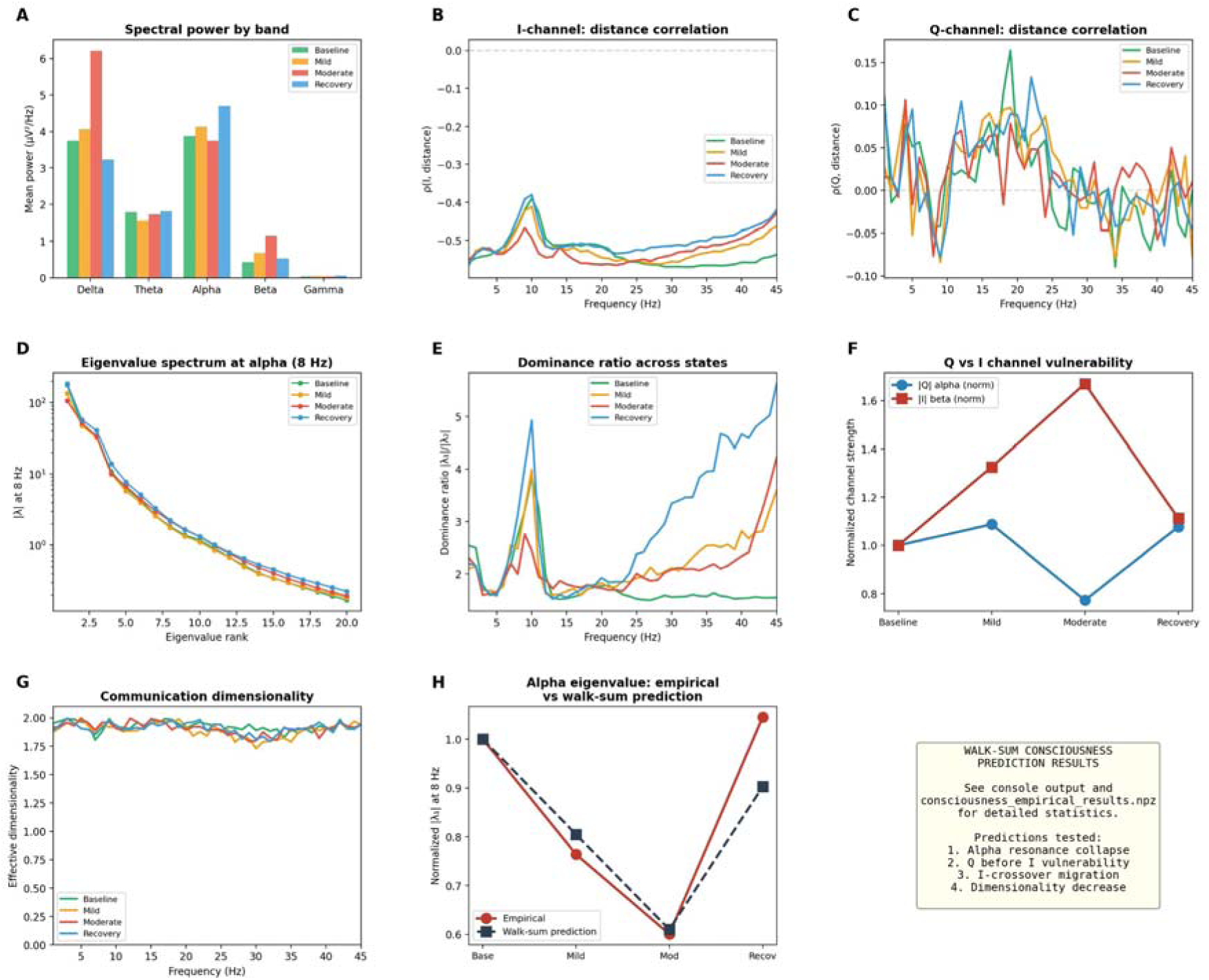
Walk-sum consciousness predictions: propofol anaesthesia (Chennu dataset, n = 20). (A) Spectral power by band across sedation states. (B) I-channel ρ(I, distance) across frequency; moderate sedation disrupts alpha. (C) Q-channel ρ(Q, distance); alpha trough deepens under mild sedation then collapses under moderate. (D) Eigenvalue spectrum at alpha (8 Hz) across states. (E) Dominance ratio |λ□|/|λ□| across frequency. (F) Q vs I channel vulnerability: |Q| alpha collapses −71% under moderate sedation (p = 0.004), |I| beta increases +366%. (G) Communication dimensionality across states. (H) Alpha eigenvalue: empirical (red) vs walk-sum prediction (blue) across sedation states; *p < 0.05, **p < 0.01.

The distinction between communication channels and dynamics is central to this work and, we argue, to the field. Neural mass models compute what signals emerge from specific biophysical parameters and inputs; the resolvent characterises what channels exist regardless of what flows through them. These are complementary but fundamentally different questions. The neural mass negative control (ρ ≈ 0.006; Fig. 7) confirms this separation empirically: the resolvent does not approximate a simulation—it describes a different object entirely. The resolvent is to brain communication what impedance is to a circuit: it constrains which frequencies can propagate and how far, without generating the signals themselves.

**Figure 7.**
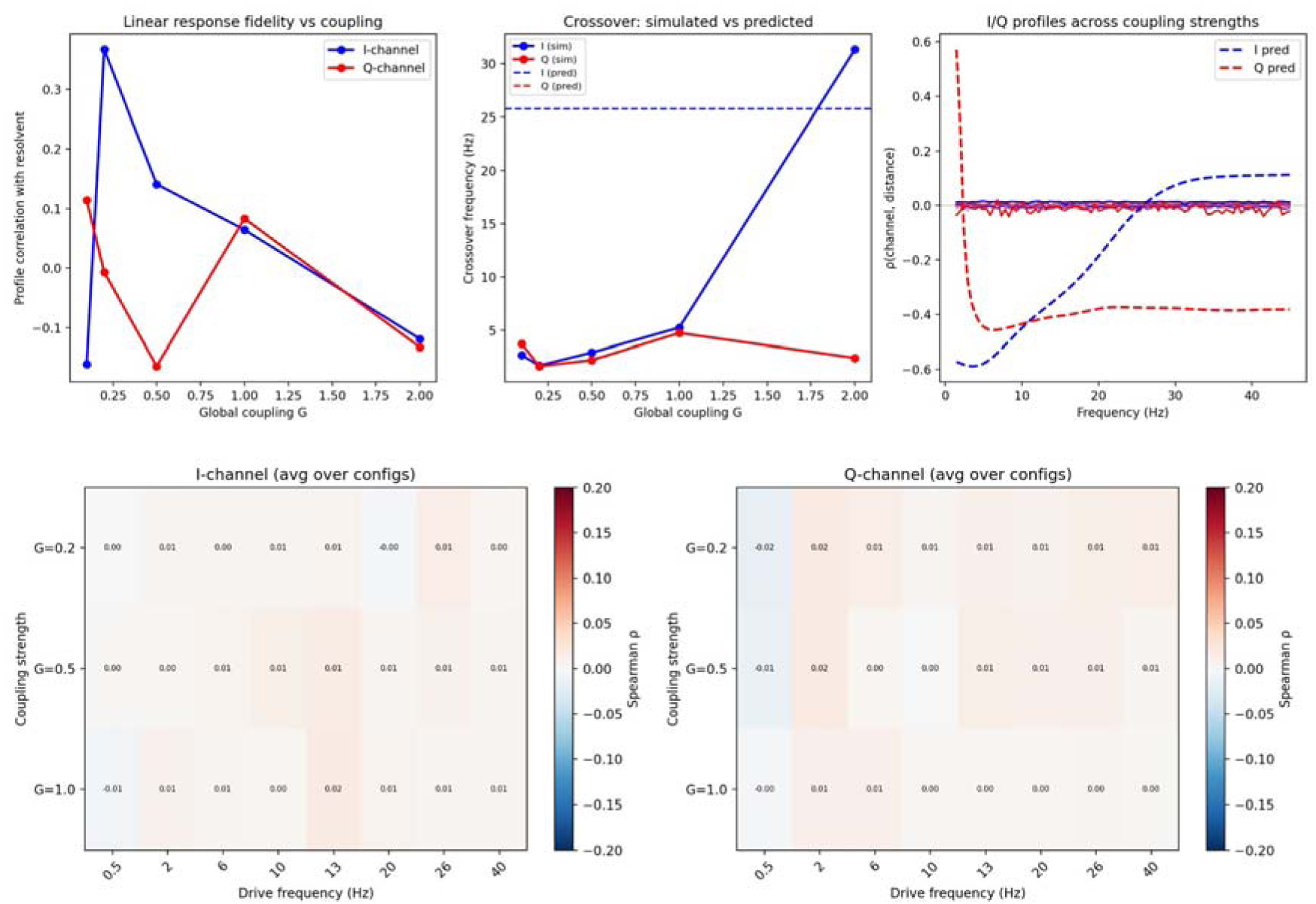
Neural mass negative control. (A) Coupling sweep summary showing spatial Spearman ρ between the resolvent’s I/Q profiles and Wilson–Cowan coherency patterns across 144 conditions (8 frequencies × 6 spatial configurations × 3 coupling strengths). Mean ρ ≈ 0.006. (B) Frequency-resolved comparison of resolvent and neural mass spatial patterns, confirming near-zero correspondence across all frequencies.

The framework’s parsimony warrants emphasis. Where neural mass models require 8–15 free parameters per node^10–15^, the bare resolvent uses none. The dressed resolvent adds exactly two—motivated not by fitting but by the biophysics of leaky neural integration^33,34^—for the entire network. That these two parameters recover the Q-crossover, improve out-of-sample prediction (Table 2), and reveal physiologically plausible hub-versus-peripheral dynamics (Fig. 4) suggests they capture genuine biology rather than statistical flexibility. The heterogeneous extension shows that hub nodes are dynamically tuned to the alpha-band corridors the topology creates, consistent with the relay physiology of thalamocortical circuits^36,37^.

Hubs and corridors play a distinctive role. High-degree nodes—predominantly in the rich club—drive the resolvent’s lowest-frequency global modes (Fig. 5A). The spatial correlation between the resolvent’s predictions and empirical connectivity is most distinct for hub-mediated communication corridors, because these corridors accumulate the most walks and therefore produce the strongest resolvent signatures. This explains why the framework’s predictions are sharpest in the alpha and beta range: these are the frequencies at which hub-mediated corridors carry the most topological structure.

The propofol and schizophrenia results illustrate the framework’s capacity to separate structural topology (**C**) from local circuit dynamics (*l*(*ω*)). Propofol alters *l*(*ω*) pharmacologically, collapsing alpha channels while preserving slower ones—a frequency-selective disruption that the dressed resolvent predicts from first principles. In schizophrenia, PV-interneuron dysfunction alters *l*(*ω*) pathophysiologically, weakening the modulatory layer and exposing the structural scaffold (topological transparency). These are not post-hoc interpretations: both follow directly from the algebraic structure of the dressed resolvent, in which **C** and *l*(*ω*) enter as separable factors. Any perturbation that changes *l*(*ω*) without changing **C** will alter the communication architecture in a manner predictable from the framework.

The framework has clear limitations. It is a linear, frequency-domain description of steady-state communication architecture. It does not capture temporal dynamics, moment-to-moment variability, or strongly nonlinear phenomena such as seizures or bursting. We maintain this is appropriate: the linear regime accounts for the 96.5% eigenmodel variance, and the channels defined by the transfer function constrain dynamics regardless of their nonlinear character. The framework also uses a single global delay *T* = 10 ms; incorporating distance-dependent delays would add biological realism at the cost of additional parameters. We have defined the falsifiability conditions for this framework—a structural connectome whose resolvent fails to predict the spatial organisation of empirical coherency—and these conditions have not been met in any of the datasets examined. Extension to time-varying regimes—in which *l*(*ω*) becomes a function of brain state—is a natural direction for future work, as are extensions to non-abelian algebraic structures that would derive cross-frequency coupling from connectome topology.

The walk-sum resolvent provides, to our knowledge, the first analytical framework that derives the frequency-dependent communication architecture of the brain from its structural connectome. By separating structural topology (**C**) from local circuit dynamics (*l*(*ω*)), it offers a principled account of how the same anatomical scaffold gives rise to different communication architectures in health, under anaesthesia (Fig. 6), and in disease.

## Methods

### Structural connectome data

Structural connectivity matrices and fibre tract distance matrices were obtained from the Bazinet *et al.* atlas [19] at four parcellation scales: Cammoun-219 (219 cortical and subcortical regions), Schaefer-400 (400 cortical regions), Schaefer-800 (800 cortical regions), and Cammoun-1000 (1,000 regions). Connectivity was derived from diffusion MRI tractography of the Human Connectome Project (HCP) young adult cohort [20], representing consensus structural connectivity averaged across a large normative sample. Matrices were normalised so that the spectral radius was below unity by dividing by 1.01 × the maximum eigenvalue magnitude when necessary. Fibre tract distance matrices represent the mean streamline length between each pair of parcels.

### MEG acquisition and source reconstruction

Resting-state magnetoencephalography (MEG) data were obtained from three independent sources. The COGITATE dataset [21] comprised 100 subjects recorded during eyes-open rest on an Elekta Neuromag 306-channel system (204 gradiometers, 102 magnetometers). The WAND dataset [22] comprised 166 subjects. The Cam-CAN dataset [44] comprised 646 subjects recorded during eyes-closed rest. Standard preprocessing included temporal signal-space separation (tSSS) for interference suppression, bandpass filtering (1–45 Hz), and artefact rejection via independent component analysis (ICA).

Source reconstruction was performed using linearly constrained minimum-variance (LCMV) beamforming [23] as implemented in MNE-Python version 1.6 [24]. Beamformer weights were computed from the data covariance matrix and the leadfield matrix derived from individual forward models based on structural MRI. Source time series were extracted at the centroids of 800 cortical parcels defined by the Schaefer atlas with 7-network assignment.

Cross-spectral density matrices were computed for each subject using Welch’s method (2,048-sample segments, 50% overlap, Hanning window) at 0.5 Hz resolution from 1 to 45 Hz. Complex coherency was computed for each parcel pair as C_{ij}(ω) = S_{ij}(ω)/sqrt{S_{ii}(ω)S_{jj}(ω)}, where S_{ij} is the cross-spectral density and S_{ii} the power spectral density.

### Intracranial EEG data and processing

The MNI Open iEEG Atlas sleep dataset [25] was obtained from Pennsieve (dataset identifier 414), comprising pre-segmented European Data Format (EDF) recordings from 108 epilepsy patients with file naming convention encoding patient, sleep stage, and epoch. After excluding 16 patients flagged for poor sleep staging and removing channels identified as resected, 90 patients were retained with data across four states: wakefulness (74 patients), N2 (89), N3 (79), and REM (57).

Electrode coordinates in MNI space were obtained from accompanying metadata. For each subject, only channels with valid MNI coordinates and not flagged as resected were included (minimum 4 channels per subject). For each subject and stage, up to 10 epochs were loaded. Data were bandpass filtered (0.5–45 Hz) and pairwise cross-spectral coherency was computed across all electrode pairs in six frequency bands: delta (0.5–4 Hz), theta (4–8 Hz), alpha (8–13 Hz), sigma (12–16 Hz), beta (16–30 Hz), and gamma (30–45 Hz). Coherency values were binned by inter-electrode Euclidean distance into 15 equal-width bins.

### DWI tractography

Diffusion-weighted imaging from the UCLA Consortium for Neuropsychiatric Phenomics (OpenNeuro ds000030) [26] was processed using DIPY version 1.9 [27]. For each of 172 subjects with DWI data: brain masking via median Otsu thresholding; diffusion tensor fitting for fractional anisotropy (FA); constrained spherical deconvolution (CSD) using auto-estimated response functions; deterministic maximum-direction tracking (maximum angle 30°, step size 0.5 mm, FA stopping threshold 0.15); 500,000 seed points in white matter; streamline-based connectivity at Schaefer-400 and Schaefer-800 parcellations. Group-average connectivity matrices were computed separately for healthy controls (n = 130) and schizophrenia patients (n = 50), symmetrised and with diagonal zeroed.

### Neural mass simulation

Noise-driven Wilson–Cowan neural mass models [28,29] were simulated on the Cammoun-219 connectome under 144 conditions (8 driving frequencies × 6 spatial input configurations × 3 coupling strengths G in {0.2, 0.5, 1.0}). Simulations used Euler integration with a 0.1 ms time step for 10 seconds per condition. Pairwise coherency was computed from simulated time series and compared to the resolvent via Spearman correlation.

### Resolvent computation and cross-validation

The bare resolvent was computed as H(ω) = (I - e^{iω T}C)^{-1} at 89 frequencies (1–45 Hz in 0.5 Hz steps) with characteristic delay T = 10 ms, using direct matrix inversion via SciPy v1.12 [30]. The dressed resolvent was computed as H_2(ω) = l(ω)(I - l(ω)e^{iω T}C)^{-1} with l(ω) = ω_0^2/(ω_0^2 - ω^2 + 2iζω_0ω). Distance-binned profiles were obtained by grouping the N(N-1)/2 upper-triangular entries into 20 equal-width bins by fibre tract distance.

Cross-validation followed a leave-one-dataset-out design: parameters (ω_0, ζ) were fitted by grid search (ω_0 in [5, 16] Hz in 0.5 Hz steps; ζ in [0.05, 1.0] in 0.05 steps) on one MEG dataset and evaluated on the other. For the heterogeneous extension, nodes were partitioned into 3 groups by weighted degree and group-specific (ω_0, ζ) were fitted via coordinate descent (3 iterations).

### Statistical analysis

The bare resolvent was computed as **H**(ω) = (**I** − e^iωT^**C**)^−1^ at 89 frequencies (1–45 Hz in 0.5 Hz steps) with characteristic delay T = 10 ms, using direct matrix inversion via SciPy v1.12^30^. The dressed resolvent was computed as **H**□(ω) = l(ω)(**I** − l(ω)e^iωT^**C**)^−1^ with l(ω) = ω□²/(ω□² − ω² + 2iζω□ω). Distance-binned profiles were obtained by grouping the N(N−1)/2 upper-triangular entries into 20 equal-width bins by fibre tract distance.

Cross-validation followed a leave-one-dataset-out design: parameters (ω□, ζ) were fitted by grid search (ω□ in [5, 16] Hz in 0.5 Hz steps; ζ in [0.05, 1.0] in 0.05 steps) on one MEG dataset and evaluated on the other. For the heterogeneous extension, nodes were partitioned into 3 groups by weighted degree and group-specific (ω□, ζ) were fitted via coordinate descent (3 iterations).

Spearman correlations were used for distance-dependent predictions; Wilcoxon signed-rank tests (α = 0.05, two-sided) for sedation comparisons. Visualisations were generated with Matplotlib^31^.

## Acknowledgements

Data collection and sharing for the Cam-CAN project was provided by the Cambridge Centre for Ageing and Neuroscience (CamCAN). CamCAN funding was provided by the UK Biotechnology and Biological Sciences Research Council (grant number BB/H008217/1), together with support from the UK Medical Research Council and University of Cambridge, UK. MEG data from the COGITATE project were provided by the COGITATE Consortium; we thank the consortium members for making these data openly available under FAIR principles. WAND MEG data were shared by Nentwich *et al.*; we gratefully acknowledge the data collection team. UCLA Consortium for Neuropsychiatric Phenomics data were obtained from the OpenNeuro database (accession number ds000030); the CNP was funded by the NIH Roadmap Initiative. Intracranial EEG data were obtained from the MNI Open iEEG Atlas via Pennsieve. Structural connectivity matrices and parcellation atlases were provided by Bazinet *et al.* and derived from diffusion MRI tractography of the Human Connectome Project. HCP data were provided by the Human Connectome Project, WU-Minn Consortium (Principal Investigators: David Van Essen and Kamil Ugurbil; 1U54MH091657), funded by the 16 NIH Institutes and Centres that support the NIH Blueprint for Neuroscience Research and by the McDonnell Centre for Systems Neuroscience at Washington University.

## Data availability

All data used in this study are publicly available. Structural connectivity matrices and parcellation atlases are available from the Bazinet *et al.* repository (https://github.com/netneurolab/bazinet_assortmix). MEG data from COGITATE are available via the COGITATE Data Release (https://www.arc-cogitate.com/data-release). WAND MEG data are available from the Parra Laboratory. Cam-CAN data are available from the CamCAN repository (https://cam-can.mrc-cbu.cam.ac.uk/dataset/). UCLA CNP data are available from OpenNeuro (accession number ds000030). Intracranial EEG data from the MNI Open iEEG Atlas are available from Pennsieve (dataset identifier 414). Propofol sedation EEG data are available from the Chennu *et al.* repository. HCP data are available from the ConnectomeDB (https://db.humanconnectome.org).

## Code availability

Analysis code is available from the corresponding author upon reasonable request.

## Author contributions

V.K. conceived the theoretical framework, wrote the first draft, performed all analyses, and curated all datasets. V.M. contributed to the interpretation of findings, helped construct the mathematical framework, and co-composed and approved the final manuscript.

